# Oral Cavity Serves as Long-Term COVID-19 Reservoir with Increased Periodontal and Viral Disease Risk

**DOI:** 10.1101/2025.02.17.638734

**Authors:** Joel Schwartz, Kristelle J. Capistrano, Raza Ali Naqvi, Sarah Elshourbagy, Armita Hezarkhani, Sodabeh Etminan, Joshua Bland, Shrihari Kadkol, Deepak Shukla, Salvador Nares, Scott L. Tomar, Afsar Naqvi

## Abstract

**Background:** SARS-CoV-2 infection can lead to long-term health problems affecting multiple body systems termed long COVID. Currently, limited information exists about long-term oral health manifestations in COVID-19 patients with limited healthcare access.

**Methods:** We conducted a sequential, cross-sectional study (December 2020–March 2024) to assess how racial/ethnic differences (Black/Hispanic vs White/Asian) and health disparities affect oral and non-oral long COVID symptoms and their relationship with COVID-19 vaccination. We retrospectively reviewed patients’ oral health record from University of Illinois Chicago dental clinics before vaccination (December 2020; N=1150; Covid+/- N=575/group) and after vaccination (December 2021; N=592; Covid+/- N=292/group). Participants were recruited in two separate prospective groups of COVID-19 positive subjects (February–April 2021; pre-vaccination: N=158; January–March 2024; post-vaccination: N=171), we examined clinical indicators of oral (periodontal and salivary glands) and non-oral (neurologic) sequalae 3–6 months after initial exposure. We measured viral S protein by flow cytometry and quantified inflammatory markers, viral entry receptors, and oral viral load to correlate molecular, and cellular changes in COVID-19 positive subjects before and after vaccination.

**Results:** Our results identified racial disparity indicating oral associated post-acute sequelae (PASC) primarily manifested as periodontal (gum) disease (COVID-19 positive: 73.1±18.9% vs COVID-19 negative: 33.1±14.3%) and correlated with higher rates of dry mouth (57.5%), taste disturbance (47%), and smell loss (20%). Vaccination reduced oral PASC in COVID-19 positive subjects; however, periodontal disease indicators persisted compared to the COVID-19 negative group. Notably, 3-6 months post-infection, while SARS-CoV-2 Spike (S) transcript was rarely detected in saliva (∼6%), its protein was commonly detected (∼70%) in the COVID-19 positive subjects indicating incomplete viral clearance. This correlates with significantly higher salivary expression of viral entry receptors (ACE2, and TRMPSS2), and inflammatory mediators (IL-6, IL-8 and MMP-8), in COVID-19 positive subjects. This finding was further supported by higher prevalence of other oral viruses including Epstein-Barr Virus (70.5%), Herpes Simplex Virus (8.1%), and Human Papillomavirus (17.5%) in COVID-19 positive subjects.

**Interpretation:** COVID-19 history significantly correlates with severe oral health complications in predominantly Black communities, while vaccination reduced but did not eliminate these issues. The oral cavity serves as a long-term viral reservoir, and periodontal inflammation with increased oral viral presence in COVID-positive patients may increase susceptibility to oral and non-oral viral diseases and identify risk for long COVID.

## Introduction

COVID-19 and its associated post-acute sequelae (PASC) impacts oral health, manifesting as xerostomia, dysgeusia, mucositis, dysphagia, sialadenitis, and mucoceles^1–3^. These oral manifestations serve as important clinical indicators of disease progression and severity. Cross-sectional and retrospective studies have identified periodontal disease (PD) as a notable pre-existing condition in COVID-19 patients, with shared inflammatory mechanisms underlying pathobiology of both diseases.^4–6^ However, the temporal evolution of COVID-19-related oral health issues and the role of pre-existing conditions like PD remains poorly understood. This knowledge gap primarily stems from limited longitudinal monitoring of oral health changes associated with SARS-CoV-2 infection, particularly when comparing the early pandemic phase (2020 to early 2021) to the post-vaccination period (mid-2021 to present). Understanding these relationships could provide crucial insights for managing both acute COVID-19 and its long-term oral health consequences.

Longitudinal studies tracking SARS-CoV-2 infection have predominantly focused on non-oral manifestations. For instance, de Miranda et al. monitored COVID-19 patients for 14 months and established that the prevalence of long COVID symptoms correlated with age, pre-existing comorbidities, and the clinical course of acute infection.^2^ Similarly, Demko et al. assessed unvaccinated individuals over 24 months and revealed that 33% experienced long COVID, with symptoms peaking at six months post-infection and significantly impacting quality of life.^3^ These investigations and more recent results from the national NIH Program-RECOVER, classified and scored for severity and prevalence various long COVID manifestations, including respiratory distress, anosmia, cognitive impairment ("brain fog"), chronic fatigue, gastrointestinal issues, myocarditis, orthostatic hypotension, neuritis, and paresthesia.^7^ However, these studies largely overlook persistent oral health complications following SARS-CoV-2 infection. Examining is the role of oral health in long COVID is particularly significant given its established connections with other chronic inflammatory conditions such as obesity, diabetes, cardiovascular diseases, osteoporosis, and chronic pain syndromes. This bidirectional relationship appears to be mediated through shared inflammatory pathways. These are initiated through a variety of phases related to SARS-CoV-2 biology: adherence, entry, with endocytosis; receptors and signals, translocation of SARS-CoV-2 virion from cytosol to nucleus and replication. This residual persistent infection is likely contributory toward oral and non-oral long COVID manifestations.

Clinical markers of periodontitis, such as increased probing depths and clinical attachment loss, and elevated systemic inflammatory markers such as C-reactive protein (CRP), ferritin, and Hemoglobin A1c (HbA1c) are insufficient to understand SARS-CoV-2 biology.^8,9^ However, dissecting the relationship between oral and systemic hyperinflammation that mediates tissue destruction may facilitate better management of long COVID to improve patient outcomes. The relationship between PD and COVID-19 has emerged as a critical area of clinical concern. Studies reveal that moderate-to-severe PD significantly increases the risk of adverse COVID-19 outcomes, including Intensive Care Unit (ICU) admission (OR=3.54, 95% CI 1.39-9.05), need for assisted mechanical ventilation (OR=4.57, 95% CI 1.19-17.4), and mortality (OR=8.81, 95% CI 1.00-77.7) compared to mild or no periodontitis^10, 11^. Equally important, though less explored, is the impact of COVID-19 on periodontal health. The presence of SARS-CoV-2 in periodontal tissues may disrupt immune homeostasis, compromising the host’s ability to control periodontal pathogens and potentially exacerbating PD^12–14^. Adding complexity to this relationship, COVID-19-induced immune dysregulation may trigger reactivation of latent oral herpesviruses, while Epstein-Barr virus (EBV) infections might contribute to both PASC development and severe COVID-19 outcomes^15–18^. Oral reservoir of SARS-CoV-2 may also be accompanied by reactivation of virion, and aid coinfections of bacterial, herpesviral and human papillomaviruses (HPV) subtypes that can be useful additional markers for long COVID.

To address these critical knowledge gaps, our comprehensive investigation combines both retrospective and prospective studies spanning 2020–2024, comparing oral health outcomes between individuals with or without COVID-19 history. Our study uniquely examines the temporal evolution of oral health changes, inflammatory status, and the presence of oral viruses—specifically SARS-CoV-2, herpesviruses, and HPV—during and after infection in both vaccinated and unvaccinated populations.

## Methods

### Human Subject Research and Consent

This study was conducted in accordance with the Declaration of Helsinki, with protocol approval from the Human Study Institutional Review Board and compliance with HIPAA guidelines to ensure patient confidentiality and rights protection (UIC IRB#2016-0696; study approval from Miles Square Research Council December 2020). These ethical approvals encompassed both the retrospective clinical observations and the collection of biological samples (saliva and gingival crevicular fluid).

*Inclusion criteria*: Adults aged 21–70 years, no oral activity (eating, rinsing, or tooth brushing) for at least one hour prior to sampling, no antibiotic use within one week of participation.

*Exclusion criteria*: Active lung disease, current COVID-19 infection.

### Study Populations

Participants in this study were selected by using probability sampling of patients in the UIC-COD electronic patient database who met the study’s inclusion criteria. First, the patient population was divided into two temporal cohorts: (i) Pre-vaccination phase (December 2020 – November 2021): Self-reported COVID-19 positive or negative subjects; and (ii) Post-vaccination phase (December 2021 to March 2024): Participants with confirmed COVID-19 history within the preceding 3–6 months, stratified by vaccination status. Non-oral pathologies were self-reported during survey and, when possible, verified with oral examination. Findings from periodontal examinations conducted in 2020–2022 were recorded in AxiUM, the electronic patient record system used at the University of Illinois Chicago College of Dentistry (UIC-COD).

We stratified our study population based on SARS-CoV-2 infection history (positive or negative). The retrospective case-control study utilized data from the UIC-COD AxiUM electronic patient database (N≈25,000), incorporating self-reported case status and clinical documentation by faculty or dental students. To ensure data integrity, we eliminated potential cross-year duplications. Our sampling began in December 2020 with documented COVID-19 infections and continued through 2024, including vaccination status data. The database provided comprehensive information on COVID-19 status, oral health parameters, and demographic characteristics including gender and race/ethnicity. Additionally, we conducted two prospective studies (2021, and 2024) in predominantly Black communities in Chicago.

### Periodontal examination

In the retrospective case-control study, trained examiners conducted full-mouth periodontal examinations of all study participants. Periodontal measurements were made at six sites per tooth (mesio-buccal, buccal, distal-buccal; mesio-lingual, lingual, and distal-lingual). All clinical assessments were performed under standardized conditions at dental clinics at UIC-COD or Mile Square, a federal qualified health center. Probing pocket depth (PPD) and clinical attachment level (CAL) assessments employed a UNC-15 periodontal probe with 1 mm calibration (HU-Friedy Mfg. Co., Inc., Chicago, IL, USA) under consistent pressure (0.20 N) for all examinations. The prospective cohort studies implemented a periodontal screening and recording (PSR) method.^19^ Periodontal indices included clinical attachment level (CAL, measured as the distance in millimeters from the cementoenamel junction to the gingival margin. Other clinical assessments of periodontal health included Plaque Index (PI)^20^, Gingival Index (GI)^21^, and Bleeding Index (BI).^22^

Gingival crevicular fluid (GCF) assessment was conducted as a measure of periodontal inflammation. The collection process involved carefully placing PerioPaper strips into the gingival sulcus, with measurements obtained by using individually calibrated Periotron 8000 devices (Oraflow Inc.).^20–23^ To ensure comprehensive evaluation of periodontal health status, multiple sampling sites were assessed for each study participant.

### Periodontal Disease Diagnostic Classifications

Periodontal diagnosis was based on established staging (stage I–IV) and grading (A, B, C) classification. ^24^

### Saliva Sampling

Subjects provided stimulated saliva samples by chewing sterile paraffin wax for 5 minutes and expectorating into a 50 ml sterile conical tube. Saliva production was evaluated for both quantity and quality. Production of less than 5 ml/5 minutes indicated xerostomia, while consistency was noted as either mucoid and stringy-viscous or watery. Saliva samples were prepared for flow cytometry and cytology analysis by aliquoting 1 ml of saliva and diluting in PBS. For RNA isolation, 1 ml of Qiazol (Qiagen, Germantown, MD, USA) was added to 1 ml of saliva and stored in -80 °C until further use. DNA was isolated from 500 μl saliva using Qiagen DNA isolation kits according to manufacturer’s instructions.

### Salivary single cell suspension and S protein detection

Salivary cells were isolated by the process optimized in our laboratory. Briefly, saliva samples were collected in 50 mL tubes and diluted 1:10 in saliva dilution buffer (SDB: 1x PBS-/- containing 2% FBS and 0.5% paraformaldehyde). The diluted samples were filtered through a 70μ filter and centrifuged at 400xg for 5 minutes. After discarding the supernatant, cells were washed 3–4 times in SDB and stored at 4°C in 2mL SDB until analysis. Flow cytometry analysis examined spike (S) protein expression both on cell surfaces and intracellularly. Isolated salivary cells (∼300,000) were washed twice with PBS containing 1% (v/v) BSA. Surface protein detection was performed with SARS-CoV-2 Spike Protein antibody (clone GT263, Invitrogen) incubation in 300 μL flow staining buffer (PBS-1% BSA) for 45 minutes on ice. Following three PBS-1% BSA washes, cells were fixed with 2% paraformaldehyde for 30 minutes at 4°C. For intracellular detection, fixed cells were washed three times with PBS-1% BSA, permeabilized with 0.3% saponin for 20 minutes at room temperature and washed again before repeating the spike antibody incubation for 45 minutes. After three washes, cells were resuspended in 300 μL PBS-1% BSA. Data from 100,000 events was acquired using a Cytek flow cytometer and analyzed with FlowJo_v10.9.0 software.

### cDNA Synthesis and Quantitative RT-PCR

Total RNA was isolated from the whole saliva of COVID(+) (n=20) and COVID(-) (n=20) subjects by using miRNeasy micro kit (Qiagen, Germantown, MD, USA). A total of 250 ng RNA was used to synthesize cDNA, which was synthesized by using the High-Capacity cDNA Reverse Transcription Kit (Applied Biosystems, Waltham, MA, USA). IL-6, TNF-α, MMP-8, MMP-13, TMPRSS2, TMPRSS4, TMPRSS13, and ACE2 (purchased from Millipore Sigma) expression levels were analyzed in a StepOne 7500 thermocycler (Applied Biosystems). The mean Ct values of three replicates were analyzed to calculate log2 fold change using the 2^−ΔΔCt^ method and normalized to β-actin.

### Nucleic acid extraction and virus analysis

Total viral nucleic acid was extracted on the EZ1 advanced extractor using the EZ1 virus mini kit v2.0 (Qiagen, cat no. 955134) according to the manufacturer’s protocol. A volume of 300µl of saliva was used to extract total nucleic acid. At the final step, nucleic acids were eluted in 60µl of elution buffer and stored at -80°C until further use. Viral analysis was performed in the CLIA/CAP-accredited molecular pathology laboratory with clinically validated assays. CMV and EBV were quantitated by real time PCR assays on an ABI Quantstudio instrument using the artus CMV RGQ MDx Kit (Qiagen, cat no. 4503245) and artus EBV RG PCR Kit (Qiagen cat no. 4501205) respectively. Standard curves were constructed using five levels of target specific plasmids included in the kits (10^1^, 10^2^, 10^3^, 10^4^, 10^5^ copies for CMV and 2.5x10^1^, 2.5x10^2^, 2.5x10^3^, 2.5x10^4^, 2.5x10^5^ copies for EBV). Each 25µl CMV PCR reaction contained 5µl of nucleic acid extract, 16.6µL CMV RG Mix and 3.4µL CMV Mg-Sol. Each 25µl EBV PCR reaction contained 5µl of nucleic acid extract, 19µL EBV PCR solution, 0.5µl EBV Probe ASR and 0.5µl EBV Primer ASR. High and low positive controls (extracts from positive patient specimens) and a no template control were included in each run. All samples, standards and controls were run in duplicate. The average Ct value from both reactions was extrapolated to the standard curve to determine the copies/reaction. Copies/reaction was then converted to copies/ml (multiplied by 40). Mean correlation co-efficient and slopes for the standard curves were 0.9953 and -3.39 for CMV and 0.9971 and -3.33 for EBV. The quantifiable ranges for CMV and EBV viral loads were 4x10^2^ to 4x10^6^ copies/ml and 10^3^ copies/ml to 10^6^ copies/ml respectively. The analytical sensitivities were 8 copies/reaction (320 copies/ml) for both CMV and EBV. Both assays were analytically specific and did not amplify other herpes viruses, common pathogens, cellular or bacterial DNA. Primers and probe sequences are proprietary to Qiagen and are not included in the package inserts.

HSV1, HSV2, VZV, SARS-CoV-2 were analyzed qualitatively by in-house developed and clinically validated real time PCR assays. HPV was analyzed by an in-house developed and clinically validated nested PCR/Sanger sequencing assay. Primers/probe sequences and assay conditions are given in **Supplementary Table X**. For HSV1, HSV2 and VZV, each 25µl reaction contained 12.5µl of 2X iTaq Universal probes supermix (Biorad, cat no. 1725130), 1 µl of forward and reverse primers (10pmols/µl), 1µl of FAM labeled probe,

4.5µl of PCR grade water and 5µl of nucleic acid extract. The analytical sensitivity of HSV1, HSV2 and VZV assays was 5 copies of target/reaction (200 copies/ml). For SARS-CoV-2, each 25µl reaction contained 6.25µl of 4X TaqPath 1-Step RT-qPCR Master Mix, CG (Thermofisher, cat no. A15299), 1µl of each primer (S, E, RNaseP genes at 10pmols/µl), 1µl of labeled probe (FAM - S, E probes at 10pmols/µl and CY5-RNaseP probe at 1pmol/µl), 1.75µl of PCR grade water and 5µl of nucleic acid extract. The analytical sensitivity of the assay was 8 copies/reaction (320 copies/ml). For HPV, analysis, each 50µl first round PCR reaction contained 5µl of 10X PCR buffer, 0.5µl of HotstarTaq DNA polymerase (Qiagen cat no. 203443), 1µl of dNTP mix (10mM, Sigma cat no. 72004), 5µl of forward and reverse primers (10pmols/µl), 28.5µl of PCR grade water and 5µl of nucleic acid extract. Each 50µl nested HPV PCR reaction contained 5µl of 10X PCR buffer, 0.5µl of HotstarTaq DNA polymerase (Qiagen cat no. 203443), 1µl of dNTP mix (10mM, Sigma cat no. 72004), 2µl of forward and reverse primers (10pmols/µl), 37.5µl of PCR grade water and 2µl of 1:5 dilution of the first round PCR product. The nested PCR product was electrophoresed on a 2% agarose gel. Reactions with a PCR product of ∼150bp were purified using the Qiaquick PCR purification kit (Qiagen cat no. cat no. 28104) with a final elution volume of 25µl. 10ng of the purified nested PCR product was Sanger sequenced bi-directionally using first round PCR primers (GP5+, GP6+). HPV genotype was determined by BLASTn alignment (https://blast.ncbi.nlm.nih.gov/Blast.cgi). Appropriate positive, negative and no template controls were included in each run for all assays. Because every saliva specimen is expected to contain cellular material, a positive RNaseP reaction in the SARS-CoV-2 assay served as an internal control to ensure successful nucleic acid extraction and rule out PCR inhibition.

### Statistical Analysis

Primary analyses included multivariate ANOVA (one-way and two-way) to determine significant associations between study groups (p<0.01). The relationship between COVID-19 infection history and poor health outcomes was assessed by using odds ratios (significance level 5%, confidence interval 95%). Between-group variance was evaluated by using F-statistics, with post-hoc analysis performed via Tukey’s Honestly Significant Difference (HSD)/Tukey-Kramer test. Effect size was quantified through linear regression (R^2^=η^2^). For COVID-19 and vaccination groups, the Kruskal-Wallis test assessed equality of variances with subsequent ranking and p-value determination.^25^

Additional analyses included multivariable linear regression to evaluate periodontal-gingival inflammation covariates, while Poisson linear modeling was used to assess probability distributions of covariant associations. Microbiome interactions and longitudinal microenvironment changes were analyzed by using Bayesian distribution models. Multiple logistic regression modeling was used to estimate odds ratios for gingival disease in the presence of multiple explanatory variables.

## Results

### COVID-19 Infection Associates with Multiple Adverse Oral Health Outcomes

We evaluated oral health changes from October 2020 through April 2024, examining COVID-19 transition from pandemic to endemic status and the influence of vaccination and booster use. **Table 1** presents our retrospective cross-sectional study populations from December 2020 (N=1150; 575 per group) and December 2021 (N=592; 296 per group). Study groups were categorized based on self-reported COVID-19 history (3-6 months post-infection) representing unvaccinated and vaccinated individuals either infected or not infected with SARS-CoV-2 during the early (testing unavailable) pandemic phase (**Table 1**). Notably, Black men and women represented the highest proportion of participants across all demographic groups (**Table 1**). The racial/ethnic composition of these communities was: Black (56.55±0.63%), White (27.3±2.12%), and Hispanic /Asian (16.15±1.48 %) (**Table 1**). Detailed racial, ethnic, and gender distributions are provided in ***Supplementary Table 1***.

**Table 1.** Demographic Characteristics of Retrospective Pre- (December 2020) and Post-Vaccination (December 2021) Case-Control Study Participants (by COVID-19 History and Race/Ethnicity).

|  | Pre-vaccination Retrospective Cohort December 2020 (N=1150) |  |  |  | Post-vaccination Retrospective Cohort December- 2021 (N=592) |  |  |  |
| --- | --- | --- | --- | --- | --- | --- | --- | --- |
| Population | n | Male | Female | Age (in years) | n | Male | Female | Age (in years) |
| History of COVID-19 | 575 | 263 | 312 | 38.5±11.22 (P= 0.202) | 296 | 133 | 163 | 46.6±22.4 (P= 0.177) |
| No History of COVID-19 | 575 | 215 | 360 | 42.0±13.51 | 296 | 108 | 188 | 44.6±17.45 |
| Population by Race/Ethnicity | Black | White/Asian | Hispanic |  | Black | White/Asian | Hispanic |  |
| COVID-19+ | 340 | 129 | 106 |  | 159 | 71 | 66 |  |
| COVID-19- | 359 | 149 | 67 |  | 177 | 69 | 46 |  |

In both pre-vaccinated and early pandemic cohort (December 2020) and post-vaccinated cohort (December 2021), COVID-positive patients showed significantly deeper PPDs compared to COVID-negative patients [(2020: COVID19+ average PPD: 5.4±1.5 mm; COVID19-average PPD: 1.9±1.0 mm; p<0.0001), (2021: COVID 19+ average PPD: 3.6±1.1 mm; COVID19-average PPD: 2.6±1.0 mm; p<0.0001)] (***Figure 1***). COVID19+ subjects also maintained over time higher values across all metrics: (1) missing teeth [(December 2020: COVID19+: 5.6±3.6; COVID19-: 2.1±1.9; p<0.0001), (December 2021: COVID19+: 5.6±3.6; COVID19-: 2.2±1.9; p<0.0001)], (2) bleeding index [(December 2020: COVID19+: 4.5±1.4; COVID19-: 1.8±1.1, p<0.0001), (December 2021: COVID19+: 3.5±0.9; COVID19-: 1.5±0.3; p<0.0001)], (3) decayed teeth [(December 2020: COVID19+: 4.2±1.2; COVID19-: 1.2±0.9; p<0.0001), (December 2021: COVID19+: 4.8±1.2; COVID19-: 2.8±0.8; p<0.0001)], and (4) fillings [(December 2020: COVID19+: 6.1±2.2; COVID19-: 4.3±1.6; p<0.0001), (December 2021: COVID19+: 6.4±0.8; COVID19-: 3.4±0.5; p<0.0001)] compared to COVID19-subjects (***Figure 1***). While these trends were consistent across genders, COVID19+ males showed higher bleeding indices than females in both cohorts: (December 2020: males: 4.8±0.4; females: 3.7±0.9) and (December 2021: males: 3.5±0.9; females: 2.7±0.4) (***Supplementary Table 1***). These results strongly suggest that SARS-CoV-2 infection may contribute to accelerated periodontal disease progression and increased susceptibility to dental pathologies.

**Fig. 1.**
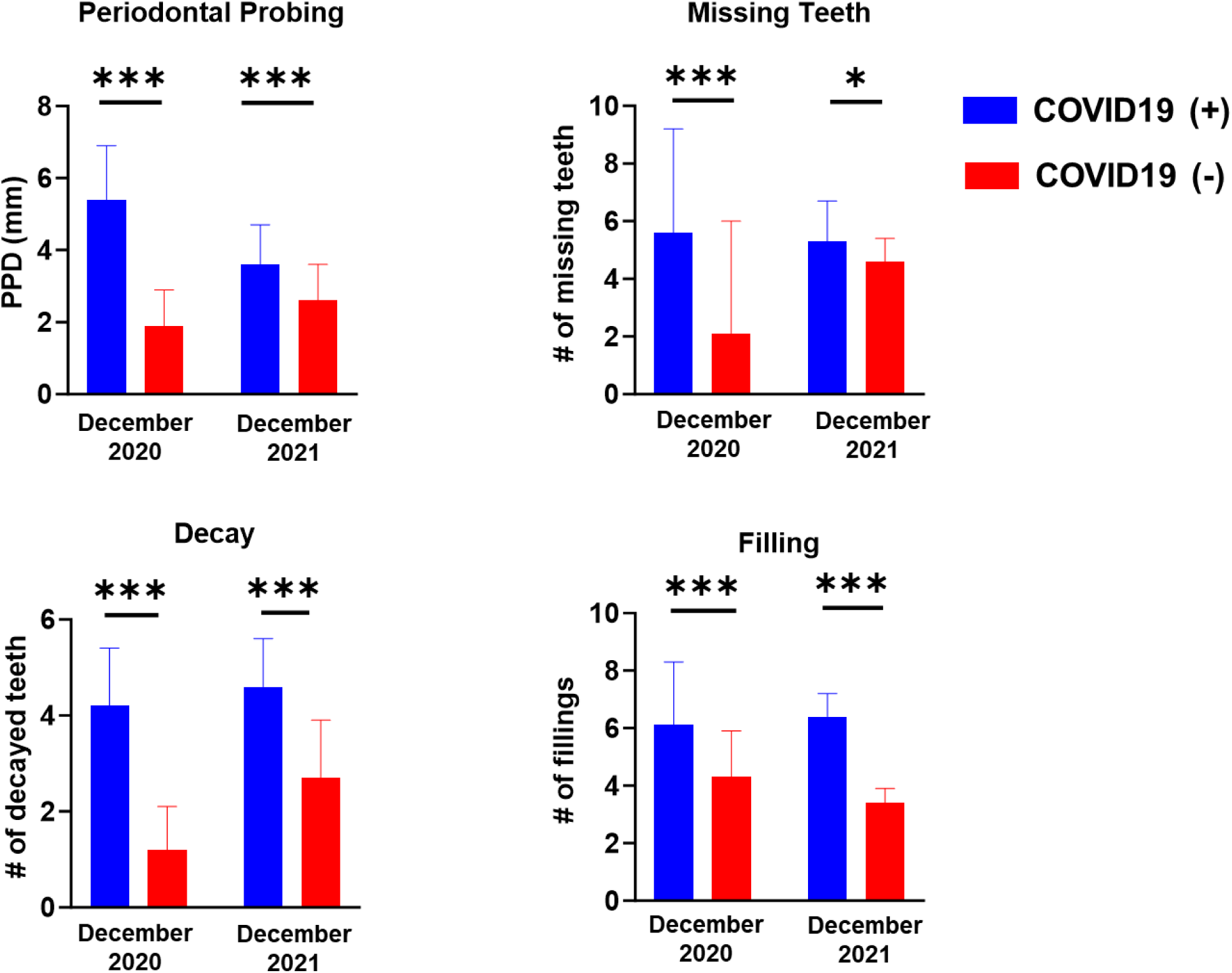
Oral Health Characteristics Related to COVID-19 History Before Widespread Vaccine Availability (Retrospective Cohorts December 2020 and December 2021).

### Vaccination Mitigates Oral Inflammatory Indices in COVID-19 subjects

These initial findings of significant oral health deterioration in COVID-19 positive individuals prompted investigation into whether vaccination could mitigate adverse oral manifestation of SARS-CoV-2 infection. To evaluate the impact of COVID-19 and vaccination on oral health, we conducted a retrospective, case-control cross-sectional study from December 2020 ─ March 2022 (N=530). Statistical analysis using ANOVA and Kruskal Wallis multiple comparisons with Dunn’s Mann-Whitney testing revealed that BI differences between groups (p=6.71e-7) were primarily influenced by COVID-19 history rather than vaccination status. COVID19+ groups showed higher BI values [COVID19+Vac+ (3.2±0.7); COVID19+Vac-(3.8±0.3)] compared to COVID19-groups

[COVID19-Vac+ (1.8±1.9); COVID19-Vac-(1.9±0.6)] (***Table 2, Supplementary Table 2***). Notably, decayed teeth counts were higher in unvaccinated groups [COVID19+Vac-(4.4±1.1); COVID19-Vac-(4.2±1.2)] versus vaccinated groups [COVID19+Vac+ (2.9±0.6); COVID19-Vac+ (2.6±0.9)]. For fillings, COVID19+ groups [COVID19+Vac+ (5.0±1.0); COVID19+Vac (3.8±0.7)] showed higher values regardless of vaccination status compared to COVID19-groups [COVID19-Vac-(3.4±2.0); COVID19-Vac+ (3.0±1.0)]. The findings indicate that while COVID-19 infection alone can contribute to worsened oral and periodontal health outcomes, while vaccination does not show any significant impact on these parameters.

**Table 2.** Retrospective Analysis of Vaccination and COVID History on Oral Health Parameters. Data was Collected from AxiUm dental records (N=530) between December 2020 ─ March 2022. *COVID+ self-reported.

| Oral Health Indices<br>(mean ± SD) | *Covid19+Vac+<br>(N=54) | *Covid19+Vac-<br>(N=23) | Covid19-Vac-<br>(N=68) | Covid19-Vac+<br>(N=385) |
| --- | --- | --- | --- | --- |
| Bleeding Index | 3.2±0.7 | 3.8±0.3 | 1.9±0.6 | 1.8±1.9 |
| Decay | 2.9±0.6 | 4.4±1.1 | 4.2±1.2 | 2.6±0.9 |
| Filling | 5.0±1.0 | 3.8±0.7 | 3.4±2.0 | 3.0±1.0 |

**Table 3.** Demographic Characteristics of Prospective Pre- (December 2021) and Post-Vaccination (January 2024) Case-Control Study Participants (by COVID-19 History).

|  | Post-vaccination Prospective Cohort December 2021 (N=84) |  |  |  |  | Post-vaccination Prospective Cohort December 2020 (N=67) |  |  |  |
| --- | --- | --- | --- | --- | --- | --- | --- | --- | --- |
| Population | n | Male | Female | Age<br>(in years) |  | n | Male | Female | Age<br>(in years) |
| History of COVID-19 | 47 | 24 | 23 | 48.0±14.09<br>(P= 0.742) |  | 44 | 15 | 29 | 46.7±14.6<br>(P= 0.727) |
| No History of COVID-19 | 37 | 12 | 25 | 49.2±19.30 |  | 23 | 10 | 13 | 45.3±16.63 |
| Population by Race/Ethnicity | Black | White/Asian | Hispanic |  |  | Black | White/Asian | Hispanic |  |
| COVID-19+ | 36 | 5 | 6 |  |  | 37 | 3 | 4 |  |
| COVID-19- | 30 | 4 | 3 |  |  | 13 | 3 | 7 |  |

### COVID-19 Exacerbate Racial Health Disparities by Impacting Oral Health Outcomes

Next, we analyzed the associations between COVID-19 history, vaccination status, and oral health across various racial and ethnic groups. Our comprehensive analysis of COVID-19 impacts on oral health revealed distinct patterns across racial, ethnic, and gender demographics. The 2020-2021 retrospective studies, which predominantly included Black cohorts, demonstrated that COVID-19+ individuals exhibited significantly compromised oral health metrics compared to COVID-19-subjects across all genders. The most striking findings emerged among Black COVID-19+ participants from 2020, who showed remarkably elevated odds ratios across all measured indices when compared to other racial groups: Bleeding Index [OR=19.6], Decayed Teeth [OR=16.4], Missing Teeth [OR=8.3], Filled Teeth [OR=6.02], and PPD [OR=5.5] (***Supplementary Table 3***). Although this pattern of deteriorated oral health persisted across Black, White, and Hispanic populations, White cohorts uniquely demonstrated non-significant increases in decayed teeth in the pre-vaccinated cohort (***Figure 2***), suggesting potential racial oral health disparities.

**Fig. 2.**
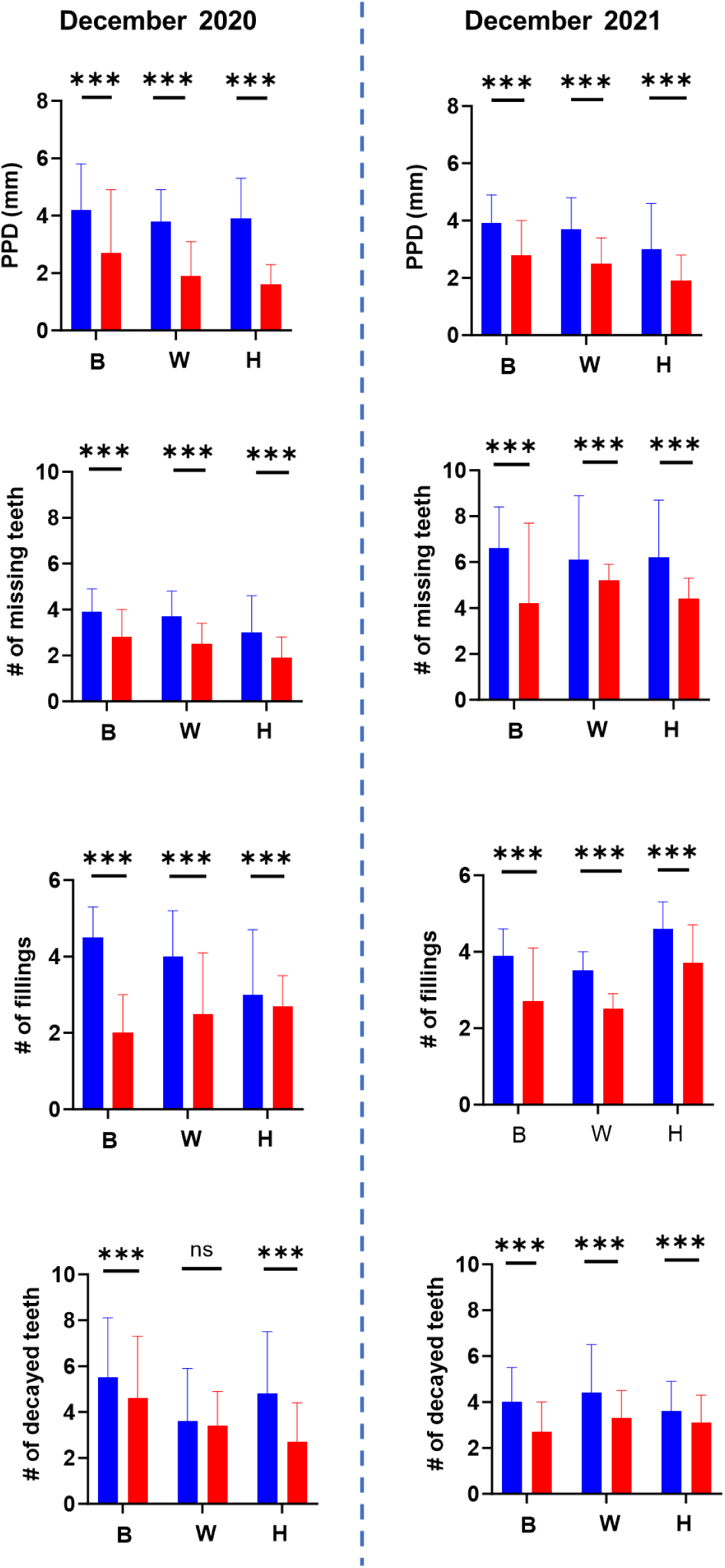
Oral Health Characteristics Related to Race/Ethnicity and COVID-19 History Before Widespread Vaccine Availability (Retrospective Cohort (December 2020 and December 2021); B=Black, W=White, H=Hispanic).

Temporal analysis spanning 2020-2021 revealed progressive improvements for all COVID+ adults, with particularly marked changes among Black participants across multiple oral health indicators: PPD (adult: p<0.001; Black: p=0.032), missing teeth (adult: p=0.16; Black: p<0.0001), BI (adult: p<0.001; Black: p=0.002), decay (adult: p<0.0001; Black: p<0.0001), and fillings (adult: p=0.02; Black: p<0.0001) (***Figure 2***). Gender-specific analysis unveiled distinct patterns, with female participants demonstrating more consistent improvements across all metrics. Male participants showed non-significant changes in several parameters, including PPD (p=0.07, with 2021 values exceeding 2020), decay (p=0.44, 2021>2020), and fillings (p=0.41), though both genders exhibited significant improvements in both missing teeth and BI measurements (p<0.0001).

The influence of COVID status (COVID+ vs COVID-) remained significant across both racial groups and genders, with distinct odds ratios emerging for different demographics (Black: OR=4.06, White: OR=2.26; male: OR=3.0, p=0.006; female: OR=4.59, p=0.032) (***Supplementary Table 3***). In contrast to gender, associations maintained their significance across racial and ethnic categories, highlighting the pervasive impact of COVID-19 on oral health. The comprehensive analysis suggests that while COVID-19 affects oral health across all demographic groups, the magnitude of impact and subsequent recovery patterns vary significantly by race and gender. Black participants demonstrated both the highest initial impact and the most substantial improvements over time, indicating potential differences in vulnerability and recovery trajectories among different demographic groups.

## Pre- and Post-Vaccination Prospective Cohorts Show Pronounced Differences in Periodontal Health Indices in COVID Subjects

Next, we performed two prospective, observational studies from 2021-2024 (N=151). The cohorts represented different vaccination phases: early pandemic (February─April, 2021; limited vaccine availability; N=84) and post-widespread vaccination period (January─March, 2024; N=67), with COVID19+Vac+ and COVID19-Vac+ groups. The prospective observational studies (N=171) maintained consistent demographic representation, with participants distributed as 65% female and 35% male. Racial composition analysis revealed a predominance of Black participants (80%±7), followed by White (7.6%±3.6), Hispanic (7.4%±10.4), and API (3.9%±1.4) populations. Throughout these investigations, COVID19+Vac+ group (73.1±18.9%) consistently exhibited elevated indicators of poor periodontal health compared to COVID19-Vac+ (33.1±14.3) groups across multiple measurement parameters (***Figure 3***). Pronounced differences was observed between COVID19+Vac+ and COVID19-Vac+ groups across multiple parameters. Periotron measurements showed significantly higher readings in COVID19+Vac+ (75.9±4.8) versus COVID19-Vac+ (64.5±2.3). This pattern extended to other periodontal indices: plaque index (COVID19+Vac+: 2.0±0.28; COVID19-Vac+: 1.0±0.3), gingival bleeding index (COVID19+Vac+: 1.7±0.2; COVID19-Vac+: 0.9±0.3), CAL/ALOSS (COVID19+Vac+: 1.9±0.7; COVID19-Vac+: 1.0 ± 0.2), and periodontal pocket depth (COVID19+Vac+: 3.3±0.7; COVID19-Vac+: 2.2±0.2) (***Figure 3***). These trends remained consistent across male and female participants (***Supplementary Table 4***).

**Fig. 3.**
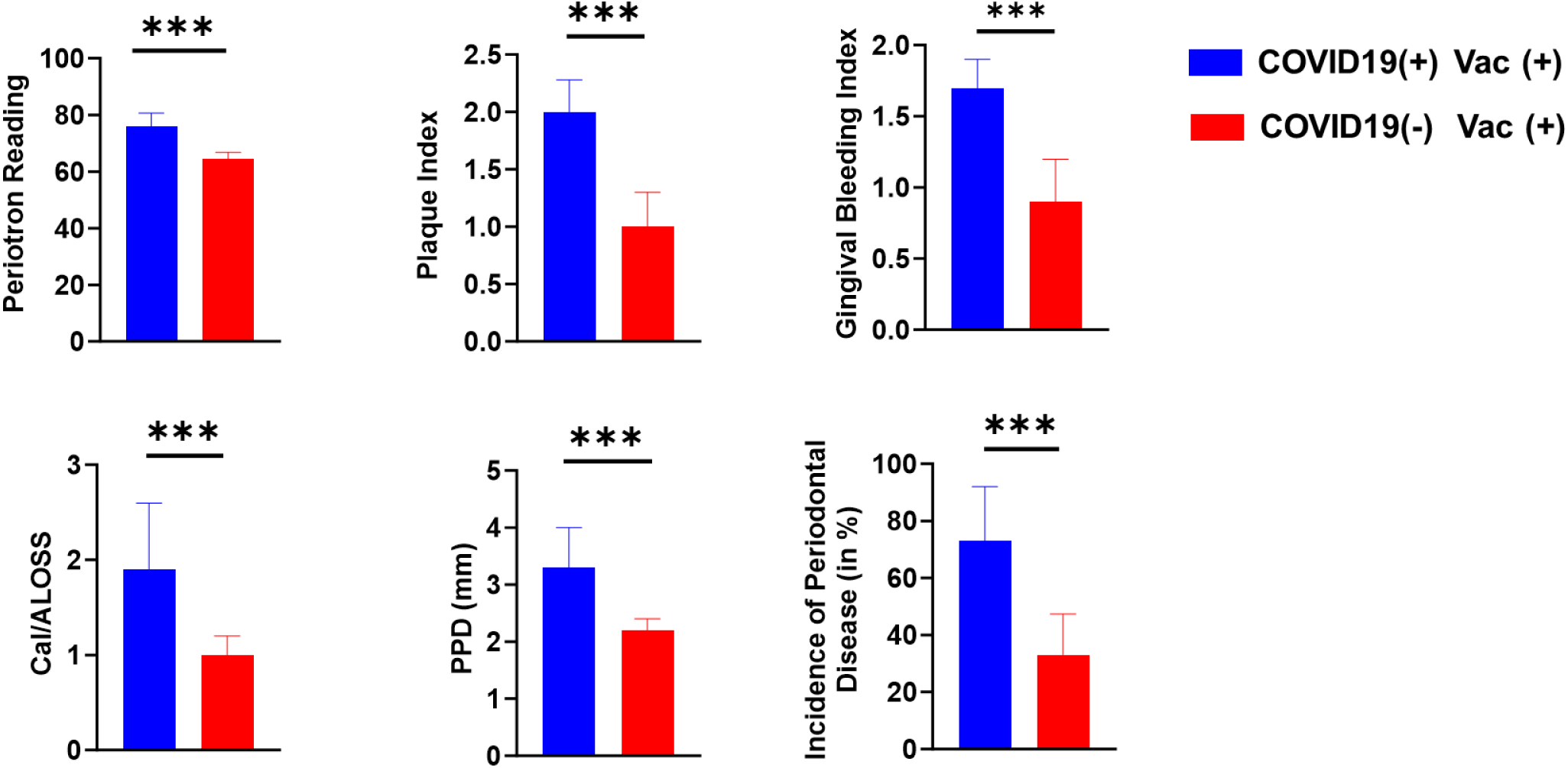
Oral Health Characteristics Related to COVID-19 Vaccination Status and History (2022-2024).

Statistical analyses uncovered a clear hierarchical pattern of significance, with CAL/ALOSS and Periotron analysis demonstrating the strongest correlations, followed by missing teeth, PI, and bleeding index measurements. Particularly noteworthy was the minimal variance observed in CAL/ALOSS and Periotron measurements among Black participants (p=0.031), while White participants showed the lowest variance in GI (p=0.011) (***Supplementary Table 4 and 5***). The post-vaccination period revealed persistent significant effects for these indexes in Black participants (p=0.002), contrasting notably with non-significant findings in White participants (p=0.380) (***Supplementary Table 4 and 5***).

### COVID-19 Subjects Exhibit Higher Prevalence of Oral and Non-Oral Long COVID Symptoms

Our previous results showed that people with a history of COVID-19 had significantly worse oral health compared to those without (COVID19-), regardless of their gender or racial background. We next examined whether oral manifestations correlate with non-oral long COVID symptoms in our prospective cohorts (N=151). Among COVID19+ patients, dry mouth (xerostomia) was extremely common, affecting 57.5% of patients, while less than 1% of COVID19-individuals experienced this condition, regardless of vaccination status, gender, or race/ethnicity (***Table 4***). Statistical analysis showed that COVID19+ individuals were >2 times more likely to develop xerostomia (p=0.022). Loss of taste (dysgeusia) was also common in COVID19+ (47.9%) compared to COVID19-(<1%) patients showing high relative risk and odds ratios (p=0.022). Loss of smell (anosmia) affected 20% of COVID19+ patients versus less than 1% of COVID19-patients reported this disorder (p=0.011) (***Table 4***).

**Table 4.** Prevalence of Oral and Non-Oral Long COVID Symptoms by COVID-19 History in Post-Vaccination Prospective Cohorts (N=151).

| **Long COVID Symptoms**<br>**(3-6 months post infection)** | **COVID19+** | **COVID19-** |
| --- | --- | --- |
| Periodontal Disease<br>(>mild gingivitis) | 73.1+/-18.9 | 33.1+/-14.3 |
| Xerostomia | 57.5% | <1% |
| Dysphagia | 47.5% | <1% |
| Anosmia | 20.0% | <1% |
| Epstein Barr Virus | 56.8% | 6.5% |
| Breathing Difficulties | ~5% | <1% |
| "Foggy Brain"-Cognitive<br>Issues | ~5% | <1% |
| Numbness-paresthesia | ~5% | <1% |
| Generalized diffuse painful<br>mouth (teeth + gingiva) | ~ 5% | <1% |
| Other Pathology | 35.5% | <1% |

The most frequent oral problem in COVID19+ patients was gum inflammation (gingivitis), affecting 73.1% of patients compared to 33.1% in COVID19-individuals. This condition, along with early-stage periodontal disease (Stages I and II), was associated with several clinical indicators: higher fluid measurements in gum pockets, increased bleeding, more plaque buildup, worse gum health scores, deeper periodontal pockets, greater attachment loss, and more missing teeth (***Table 4***). Our analysis revealed dramatically higher risks in COVID19+ patients for dry mouth (p=0.0093), difficulty swallowing (p=0.011), and loss of smell (p=0.031). Consistent with this, we noted that subjects with severe PD displayed a more viscous, mucoid, “stringy” saliva compared to more watery saliva noted for healthy or mild periodontal conditions. Other oral conditions, though less common, included oral tissue inflammation (mucositis), salivary gland inflammation (sialolitis), and fluid-filled sacs (mucocele), suggesting widespread inflammation affecting both gums and salivary tissues (p=0.016). Beyond oral symptoms, patients also reported digestive problems including cramps, bloating, and intestinal pain (35.5%). Less common symptoms in COVID19+ patients included breathing difficulties, cognitive problems, numbness/tingling sensations, and general mouth pain, each affecting approximately 5% of patients.

### Salivary Detection of S-protein Indicates Viral Persistence in Long COVID Subjects

We next examined the persistence of SARS-CoV-2 in oral tissues to understand its relationship with long COVID oral symptoms. SARS-CoV-2 presence was examined by quantifying spike transcript expression in the saliva samples collected between 2022-2024. RT-qPCR analysis revealed viral transcripts (S protein) in 6.4% of the saliva samples (***Table 5***). This is still a remarkable finding considering these samples were collected several months after the initial infection. To further confirm viral persistence, we conducted flow cytometric analysis, which revealed S-protein staining in saliva was significantly higher in COVID+ individuals in both pre-vaccination (3.2±0.91 %; p<0.001) and post-vaccination (2.9±1.12 %; p<0.001) cohorts compared to healthy individuals with no history of COVID (0.46±0.12%) (***Figure 4A,B***). Intriguingly, S-protein was also detected at higher rate (2.3±0.46%; p<0.01) in periodontally healthy individuals within one year of SARS-CoV2 infection (at the time of recruitment). Detection of S-protein irrespective of vaccine and long COVID symptoms suggests long-term persistence of virus in the oral cavity. These findings provide a potential mechanistic explanation for the prolonged oral health complications frequently observed in long COVID patients and highlight the importance of ongoing oral health monitoring in COVID-19 recovery.

**Fig. 4.**
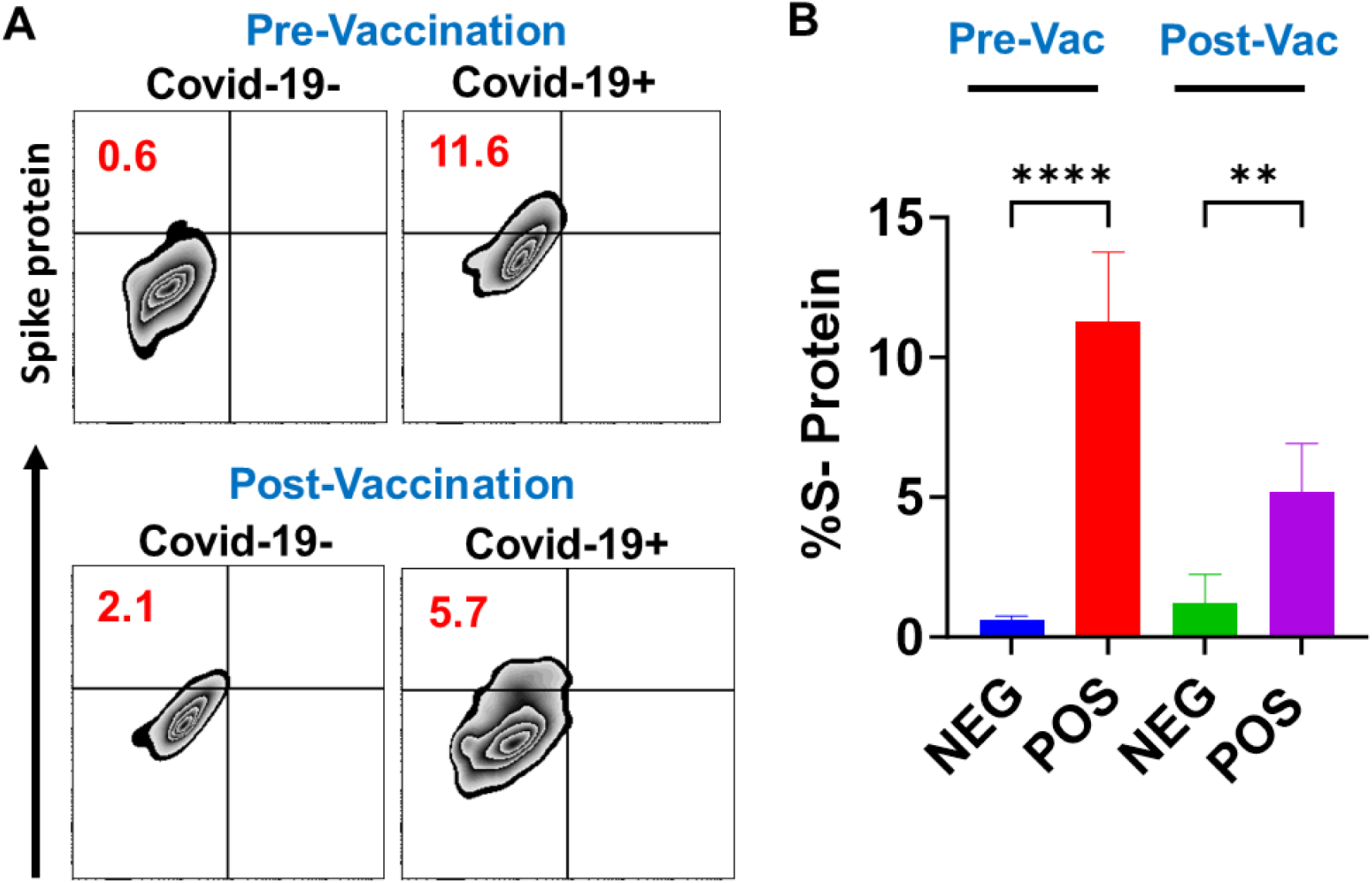
Flow Cytometric Analysis of SARS-CoV-2 S Protein Detection in the Salivary Cells of COVID-19+ and COVID-19-subjects in Pre- and Post-Vaccinated Cohorts.

**Table 5.** Prevalence of Oral Viral Infections Related to COVID-19 History. Pre- and Post-Vaccinated Prospective Cohorts (N=151) Were Pooled to Evaluate Overall Changes in Different viruses. HPV genotyping Identified Following Subtypes: HPV **3,**10, 44, 55, **56,** 72 (**bold**: High Risk Subtypes).

| Virus | Total Population (%) | COVID-19+ (%) | COVID-19- (%) |
| --- | --- | --- | --- |
| SARS-CoV-2 | 6.4 | 3.7 | 2.7 |
| EBV | 70.6 | 54.4 | 16.2 |
| CMV | 3.1 | 2.1 | 1.0 |
| HSV-1 | 8.1 | 8.0 | 0.1 |
| HSV-2 | 0 | 0 | 0 |
| VZV | 0 | 0 | 0 |
| HPV* | 17.5 | 12.5 | 5 |

### Higher Expression of Viral Entry Receptors and Immune Mediators in Long COVID

Expression of viral entry receptors in the oral cavity, combined with local immune responses, may create a favorable environment for viral persistence and chronic inflammation. This mechanism may contribute to the development and progression of oral diseases, particularly periodontal conditions, in COVID-19 patients. We therefore investigated the molecular relationship between COVID-19 and periodontal inflammation by examining immune mediators and viral entry receptors. We have previously shown higher ACE2 and IL-6 expression in inflamed gingival biopsies collected in a pre-pandemic cohort compared to healthy controls suggesting a relationship between periodontal inflammation and viral entry receptors^26^. In this pursuit, we examined salivary expression of various pro-inflammatory cytokines (IL-6 and TNF-α) and matrix metalloproteinase 8 (MMP8) that are associated with periodontal disease pathology. Our results show that the expression of IL-6 was significantly increased in COVID-19+ subjects pre- (14.95±5.29 fold; P<0.0001) and post-(7.21±2.94 fold; P<0.0002) vaccination compared to COVID-19-patients ***(Fig. 5A)***. TNF-α also showed higher expression in COVID-19+ both pre- (13.51±4.52 fold; P<0.0001) and post- (5.12±4.06 fold; P<0.013) vaccination ***(Fig. 5B).*** Similarly, higher expression of MMP8, a key periodontal tissue destructing enzyme, was observed in COVID-19+ (pre-vaccination: 44.12±29.35 fold; P<0.0001; post-vaccination: 17.55±6.90; P<0.03) subjects regardless of the vaccination status ***(Fig. 5C).*** Importantly, we noted that post-vaccination COVID-19+ subjects showed relatively less inflammatory marker levels. These findings corroborate with the previously stated poorer oral health status of COVID+ compared to COVID- and also consistent with demographic results for oral health status, with NHB/H patients with a COVID+ history even following vaccination at a higher level of poor oral health compared to other racial groups.

**Fig. 5.**
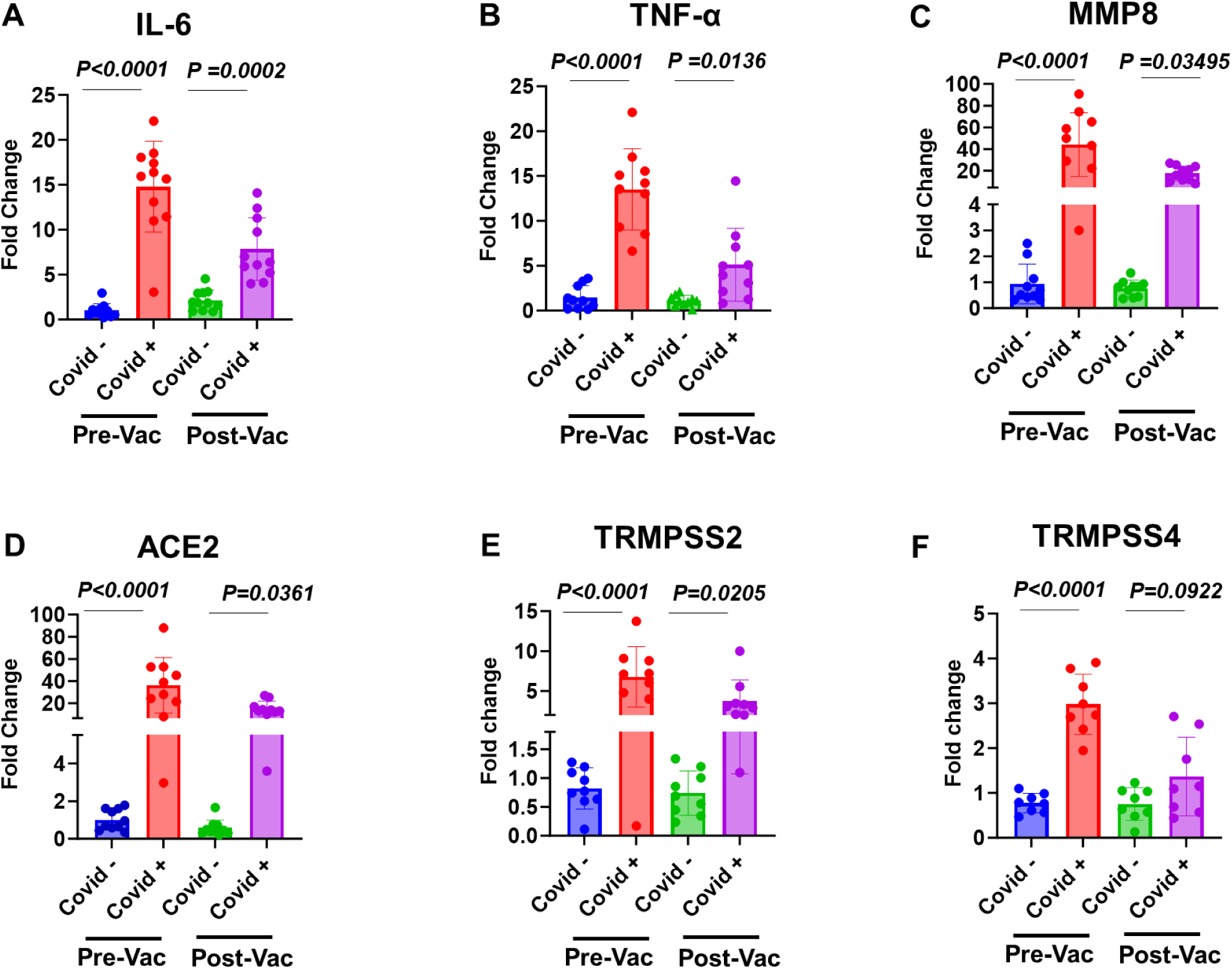
Salivary Expression Levels of Pro-inflammatory Mediators and Viral Entry Receptors in COVID-19+ and COVID-19-subjects in Pre- and Post-Vaccinated Cohorts.

Next, we evaluated the salivary expression of SARS-CoV-2 entry receptors (ACE2, TRMPSS2 and TRMPSS4) in pre- and post-vaccination cohorts. Our results show significantly increased expression of ACE2 in COVID-19+ subjects in both pre-vaccination (34.12±27.35; p<0.0001) and post-vaccination (14.9±6.88 fold; P<0.036) cohorts compared to COVID-19-individuals (***Fig. 5D***). Similarly, the expression of TRMPSS2, other entry receptor, was also significantly higher in COVID-19+ subjects regardless of vaccination status (pre-vaccination: 6.79±3.5-fold, P<0.0001; post-vaccination: 3.76±2.50 fold, P<0.02) (***Fig. 5E***). However, the levels of TRMPSS4, another SARS-CoV-2 entry receptor, were significantly higher (2.9±0.65; P<0.0001) in pre-vaccination but not in post-vaccination (1.61±0.93; P<0.09) COVID-19+ subjects (***Fig. 5F***). Overall, these findings suggest that higher inflammatory burden correlates with increased viral entry genes expression in COVID-19+ subjects suggesting a conducive microenvironment for SARS-CoV-2 infection and persistence.

### Oral Burden of DNA viruses is higher in long COVID subjects

Previous studies have demonstrated COVID-19-driven immune dysregulation may reactivate latent herpesvirus infections in the oral cavity ^27–29^. Conversely, EBV infections may be a contributing factor to the formation of PASC and severe COVID-19^15–18, 30–32^. We contend that the interplay between herpesviruses and SARS-CoV-2, while remaining largely uninvestigated, is important to decipher given the established associations between viral infections, systemic health, and oral inflammatory diseases. We quantified various herpesviruses (EBV, CMV, HSV-1, VZV and HSV-2) in saliva of pre- and post-vaccinated cohorts. Our results show that EBV was highly prevalent (70.6%) in our cohort of 2021-2024 (N=151) subjects with levels increasing with COVID-19 history (COVID19+: 54.5%; COVID19-: 16.2%) (**Table 5)**. Herpes simplex virus-1 (HSV-1) also increased in COVID19+ subjects relative to other herpesviruses (i.e., CMV, HSV2, and VZV) (**Table 5**).

Recent studies have indicated higher rates of cancers post-COVID-19 pandemic; however, the underlying etiology remains largely unknown. Human Papilloma Virus (HPV) is a unique oral oncogenic virus. We therefore investigated HPV burden by genotyping different strains and noticed elevated detection (17.2%) amongst our 2022-2023 cohort. In this predominantly Black cohort, we detected all HPV subtypes except for subtype 16 (**Table 5**). Among these, high-risk oncogenic subtypes (HPV-3 and HPV-56) were detected at higher rates in our pre-vaccinated cohort. After vaccination, our results identified higher prevalence (8.1%) of four additional high-risk subtypes (HPV-18, HPV-35, HPV-45, and HPV-56) in COVID+ subjects (**Table 5**).

## Discussion

The long-term impact of COVID-19 manifestations in the oral cavity has broad and current interest. Cross-sectional studies and case reports suggest that subjects with oral disease are more susceptible to SARS-CoV-2 infection or severity of COVID-19 symptoms.^33, 34^ Individuals from communities of poor oral health access are at risk for PASC from COVID-19.^35, 36^ However, there is very limited knowledge how long COVID impacted oral health. This study provides the first long-term analysis (2020-2024) of the impact of long COVID on oral health, combining both case-control retrospective and prospective approaches to examine the relationship between SARS-CoV-2 infection and oral diseases. By analyzing data from early pandemic stages through post-vaccination periods, we uncovered several key patterns. Across all study years, COVID19+ participants showed consistently higher rates of oral health problems compared to COVID19-individuals, including gingivitis, periodontitis, tooth decay, missing teeth, and various salivary disorders (xerostomia, dysgeusia, dysphagia, and parosmia/anosmia). These differences persisted regardless of gender, race/ethnicity, and vaccination status. These results strongly suggest that SARS-CoV-2 infection may contribute to accelerated periodontal disease progression and increased susceptibility to dental and other oral pathologies. These findings extend the pattern of susceptibility to SARS-CoV-2 infection based upon previous reports that prior immunosuppression (e.g., HIV) was present to aid infection^37^.The consistent pattern of deteriorated oral health metrics in COVID-19 positive individuals highlights the importance of enhanced dental monitoring and preventive care in patients with COVID-19 history, particularly during the early stages of infection and recovery. Clinically, increased oral health monitoring and care may mitigate long COVID symptoms, especially in underserved communities with limited oral health access.

Interestingly, we observed encouraging improvements in oral health metrics over time, particularly among vaccinated individuals. For instance, the mean PPD decreased by ∼two-folds in COVID19+ patients from 2020 (5.4 ± 1.5 mm) cohort compared to COVID19+Vac+ individuals from 2022-2024 cohort (to 3.3 ± 0.4). Similarly, the bleeding index (BI) showed improvement, dropping from 4.5 ± 1.4 in early COVID19+ cases (2020) to 3.2 ± 0.7 in vaccinated individuals in 2022-2023. These findings suggest that vaccination may play a protective role in oral health outcomes for individuals with previous COVID-19 infection. However, because this was not found in all individuals, we suggest the presence of a subset of individuals of poor responders requiring further study as they may represent a core group that eventually develop long COVID.

Our study uniquely focuses on urban Black communities, a population historically underrepresented in COVID-19 oral health research. It is also important to consider the possibility that limitations related to access to oral health care impact the higher prevalence of periodontal disease at initial exposure to SARS-CoV-2 and beginning of COVID-19 (2020) for many individuals we later examined prospectively (2023-2024). This demographic focus may explain our higher reported rates of oral manifestations. For example, we found dysgeusia in 47.5% of participants, substantially higher than the 1.6% reported in previous studies^38^. While COVID-19 impacted oral health across all demographics studied here, our early pandemic data revealed that Black participants faced significantly higher risks of poor oral health outcomes. Additionally, Black individuals showed increased susceptibility to long COVID related oral health complications, though vaccination notably reduced these risks. These findings highlight a concerning pattern where minority (Black and Hispanic) participants experienced disproportionately worse oral health outcomes compared to other demographic groups.

SARS-CoV-2 is routinely detected in saliva, and ACE2/TMPRSS expressions identify with a wide expression of gingiva, tongue, and salivary gland epithelium suggesting viral tropism for oral tissues^39^.Furthermore, the expression of viral entry receptors has been shown to correlate with inflammatory status, and both COVID-19 and PD are inflammatory, infectious diseases that involves overt immune activation.^19, 20^ Subjects with oral disease exhibit susceptibility towards viral infection; however, whether they display unique features required for viral entry, and persistence remain largely unknown. Our results demonstrate significantly higher expression of both viral receptors as well as pro-inflammatory cytokines including IL-6, TNF-α and MMP8 in COVID-19+ subjects suggesting a molecular relationship between oral inflammation and viral tropism. These molecular interactions may explain the high prevalence of oral manifestations in COVID-19 patients and their persistence in long COVID cases. Importantly, we noticed that while vaccinated COVID-19+ subjects still showed higher burden of inflammatory mediators, this burden was comparatively reduced compared to subjects in the pre-vaccinated COVID-19+ cohort.

COVID-19-driven oral pathologies may arise, in part, (1) due to the low-level persistence of SARS-CoV-2 and (2) viral-viral interactions in the oral cavity. Past studies have implicated the oral cavity as a reservoir for viral infections, including SARS-CoV-2 and herpesviruses, serving as a primary site for viral contact, entry, and replication^39, 40^ While our qPCR analysis of saliva detected SARS-CoV-2 RNA only in 6.4% of the 2022 to 2024 cohort, we detected elevated levels of salivary spike protein, ACE2, and TMPRSS2 in COVID-19+ individuals, indicating the presence of a low-level SARS-CoV-2 reservoir in the oral cavity months after primary infection. Because saliva continuously bathes all epithelial-lined surfaces of the oral mucosa, the maintained salivary expression of S-protein, ACE2, and TMPRSS2 reflects active viral replication in the oral cavity, suggesting a sustained viral load in COVID19+ individuals as the SARS-CoV-2 variants evolved from the 2019 wild type to the current 2024 Omicron variants (KP.2, KP.3, and LB.1).^41^ Further research is needed to determine which variants contribute to the persistence of the virus and impedes clearance from the oral microenvironment, potentially leading to oral long COVID manifestations^42^. We further noted increased identification of both EBV and HPV oncogenic subtypes which also suggestion possible reactivation or increased susceptibility to these oral oncogenic viruses. Further studies are needed to determine how significant this finding relates to cancer control and risk for oral carcinomas.

Our study combines clinical oral examinations with commonly detected oral DNA viruses in saliva samples, revealing increased viral loads in COVID19+ individuals, particularly for EBV, HSV-1, and HPV. This finding correlates with our observations of higher oral inflammation and diverse inflammatory salivary immune cell populations in COVID19+ subjects (*Naqvi et al. Unpublished results*). Such immune dysregulation may increase susceptibility to disease and trigger reactivation of latent herpesvirus infections in the oral cavity^43^. While herpesvirus reactivation has been documented throughout the body, studies specific for the oral cavity remain limited^44–46^. Gold et al. found EBV reactivation in 66.7% of long COVID-19 subjects, suggesting a link between COVID-19 inflammation and persistent symptoms^47^. Our findings corroborate with other research showing EBV reactivation in COVID-19 patients, particularly in long COVID cases. Additionally, we detected HPV6 oncogenic subtypes (2022-2024 cohort), which could increase risk for various carcinomas and oral papillomas. While papillomas are benign, they can compromise mucosal protection, potentially increasing infection, and inflammation risk. HPV oncogenic subtypes will express oncoproteins E5,E6,E7) depressing checkpoint cell cycle control from P53, and pRb, Cyclin D1/p27kip1, p21WAF1/ CIP1, p16INK4A. In addition, tumor suppressors genes (p53, pRB) are hypermethylated, while hypomethylation of viral genome and histone leading to alterations in noncoding RNAs (including microRNAs, circular RNAs, and long noncoding RNAs)^48^.This environment may disrupt cell cycle regulation in oral keratinocytes, promoting neoplasia development^49^. These findings emphasize the importance of investigating how SARS-CoV-2 and oral viral interactions affect immune regulation, potentially driving chronic inflammation and viral reactivation, which may contribute to long COVID and systemic immune dysfunction.

While our study provides valuable insights into oral and systemic manifestations of long covid, certain methodological considerations warrant acknowledgment. The nature of data collection relies partly on patient recollection and self-reported symptoms. The involvement of multiple clinical assessors, though standardized in approach, introduces subtle variability inherent to retrospective analyses. The snapshot nature of our single-visit design, while comprehensive, presents an inherent constraint in capturing the dynamic nature of disease progression. Additionally, biological sample collection and processing, subject to practical constraints of clinical settings, may introduce minor temporal and quantitative variations that merit consideration when interpreting results. Nonetheless, our findings reveal novel insights on SARS-CoV-2 and vaccination impact on oral health in long COVID subjects and a risk of increased viral burden on long-term chronic diseases.

## Summary

Our longitudinal study (2020-2024) demonstrates three key findings about oral health manifestation of long COVID. First, individuals with a history of SARS-CoV-2 infection showed significantly poorer oral health outcomes and higher rates of both oral and non-oral sequelae. Second, these oral health problems correlate with higher expression of inflammatory mediators in COVID19+ subjects compared to COVID19-individuals. Third, we found that oral health recovery appears to be hindered by the continued presence of SARS-CoV-2 and the presence of other oral viral pathogens. This provides a conducive microenvironment that promotes both SARS-CoV-2 persistence and ongoing inflammation in oral tissues. Targeted policy interventions and resource allocation for racial minorities are urgently needed to not only mitigate long COVID but prevent other systemic chronic diseases, which may stem from higher rates of untreated oral diseases.

## Notes

### Competing Interest Statement

The authors have declared no competing interest.

