## Supplementary Files for "Oral Cavity Serves as Long-Term COVID-19 Reservoir with Increased Periodontal and Viral Disease Risk"

**Supplementary Tables**

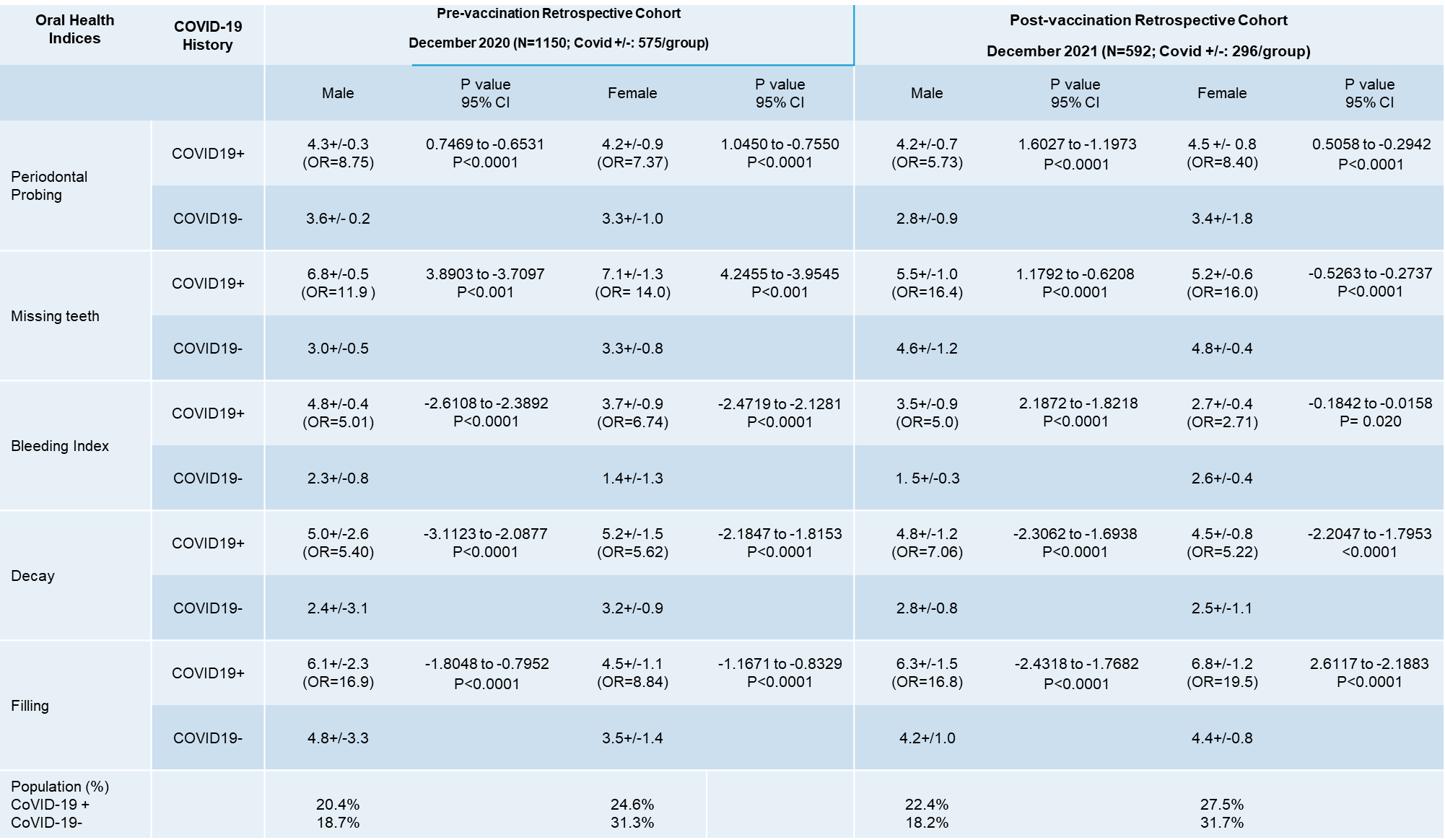

**Supplementary Table 1.** Oral Health Characteristics Related to COVID-19 History in Pre- (December 2020) and Post-Vaccination (December 2021) Retrospective Cohorts.

| **Adult** | **Covid19+Vac+**  **(N=54)** | **Covid19+Vac-**  **(N=23)** | **Covid19-Vac-**  **(N=68)** | **Covid19-Vac+**  **(N=385)** |
| --- | --- | --- | --- | --- |
| Male | 37.0 | 52.1 | 45.5 | 38.5 |
| Female | 62.9 | 47.8 | 54.4 | 61.5 |
| Black | 24.0 | 8.6 | 0.0 | 38.7 |
| White | 72.2 | 47.8 | 29.4 | 57.6 |
| Hispanic/API | 57.4 | 43.4 | 70.5 | 39.5 |

**Supplementary Table 2**. Demographics of Retrospective Cohort (December 2020 – March 2022)

to Assess the Impact of Vaccination and COVID history on Oral Health Parameters.

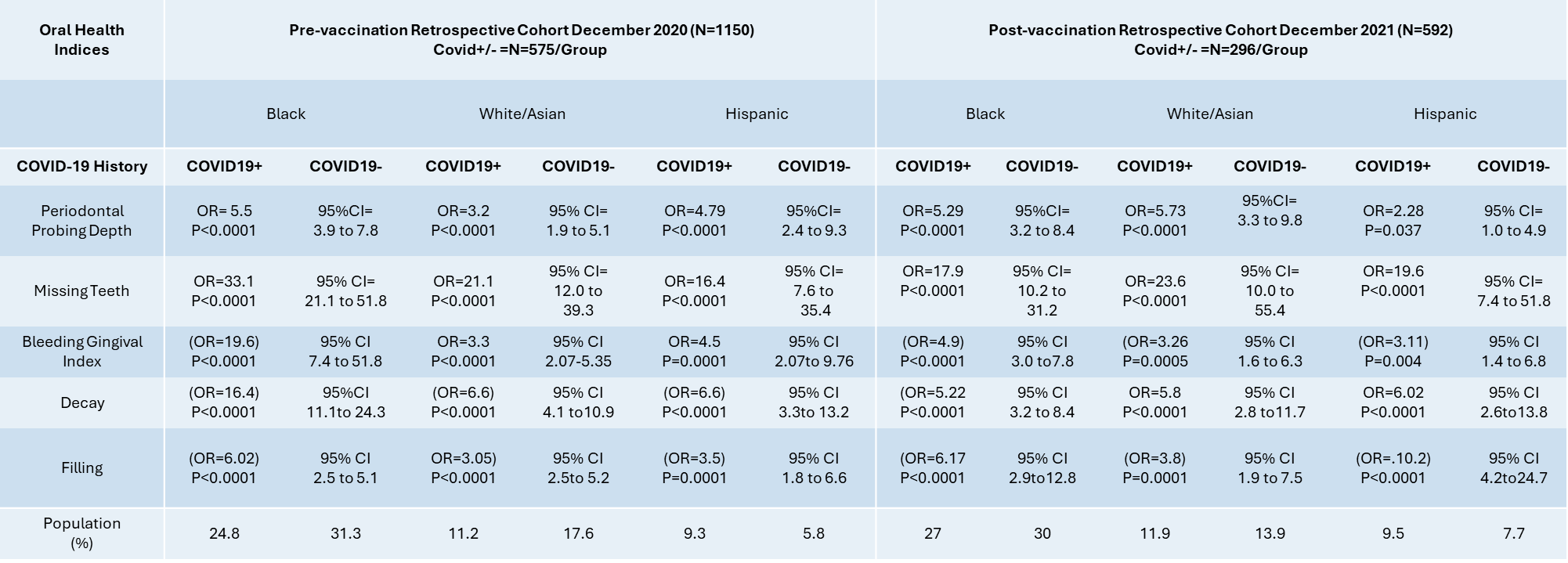

**Supplementary Table 3.** Oral Health Characteristics Related to Race/Ethnicity and Covid-19 History Before Widespread Vaccine Availability (Retrospective Cohort (2019 and 2020); B=Black, W=White, H=Hispanic) Presented as Odds Ratios (OR) with 95% Confidence Intervals (CI).

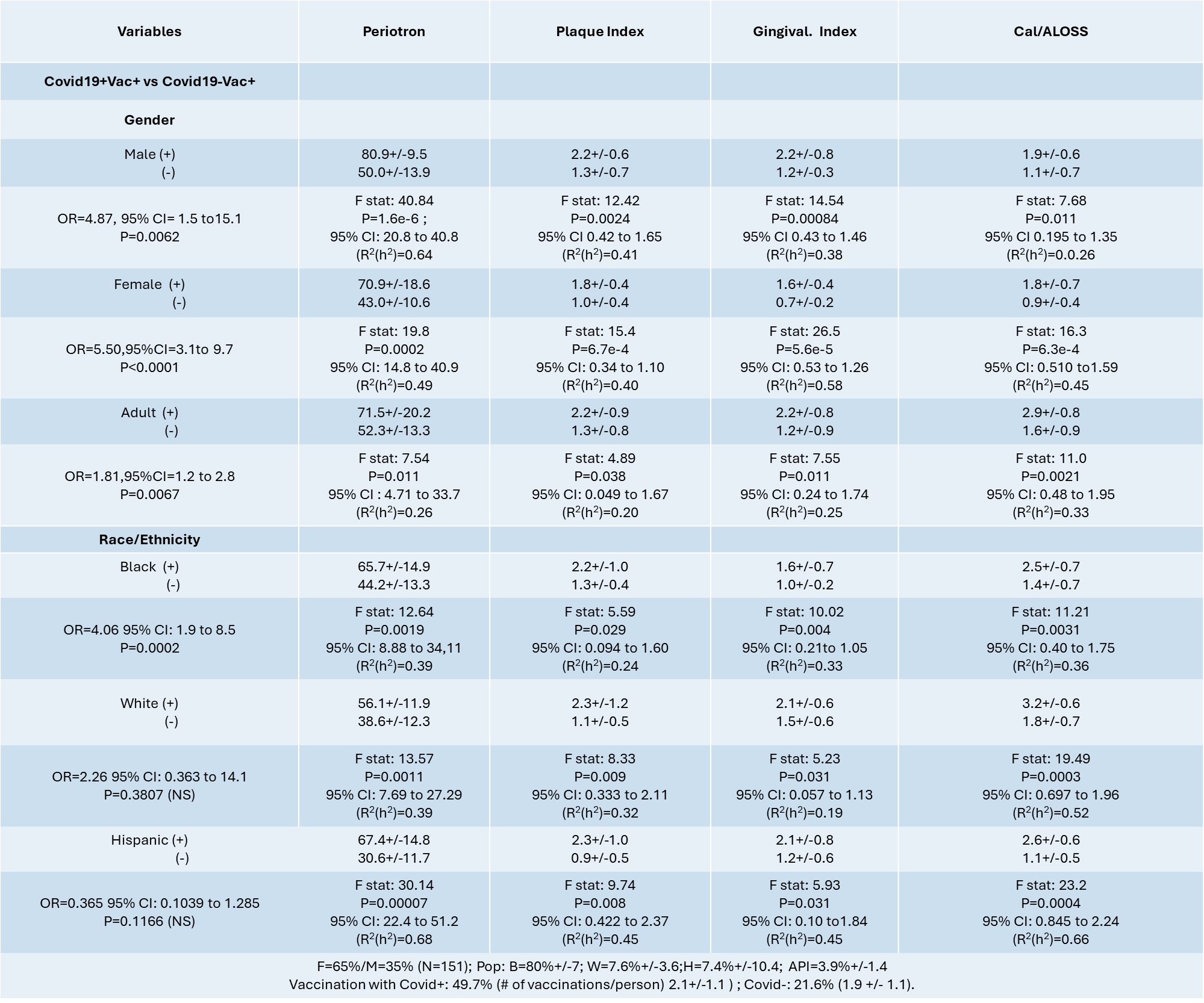

**Supplementary Table 4.** Comparative Prospective Examination Results for Oral Health Indices and Periotron Readings Associated with COVID and Vaccination (December 2021 – March 2024; N=151). Data was Stratified to Reflect Racial Differences in the Oral Health Indices.

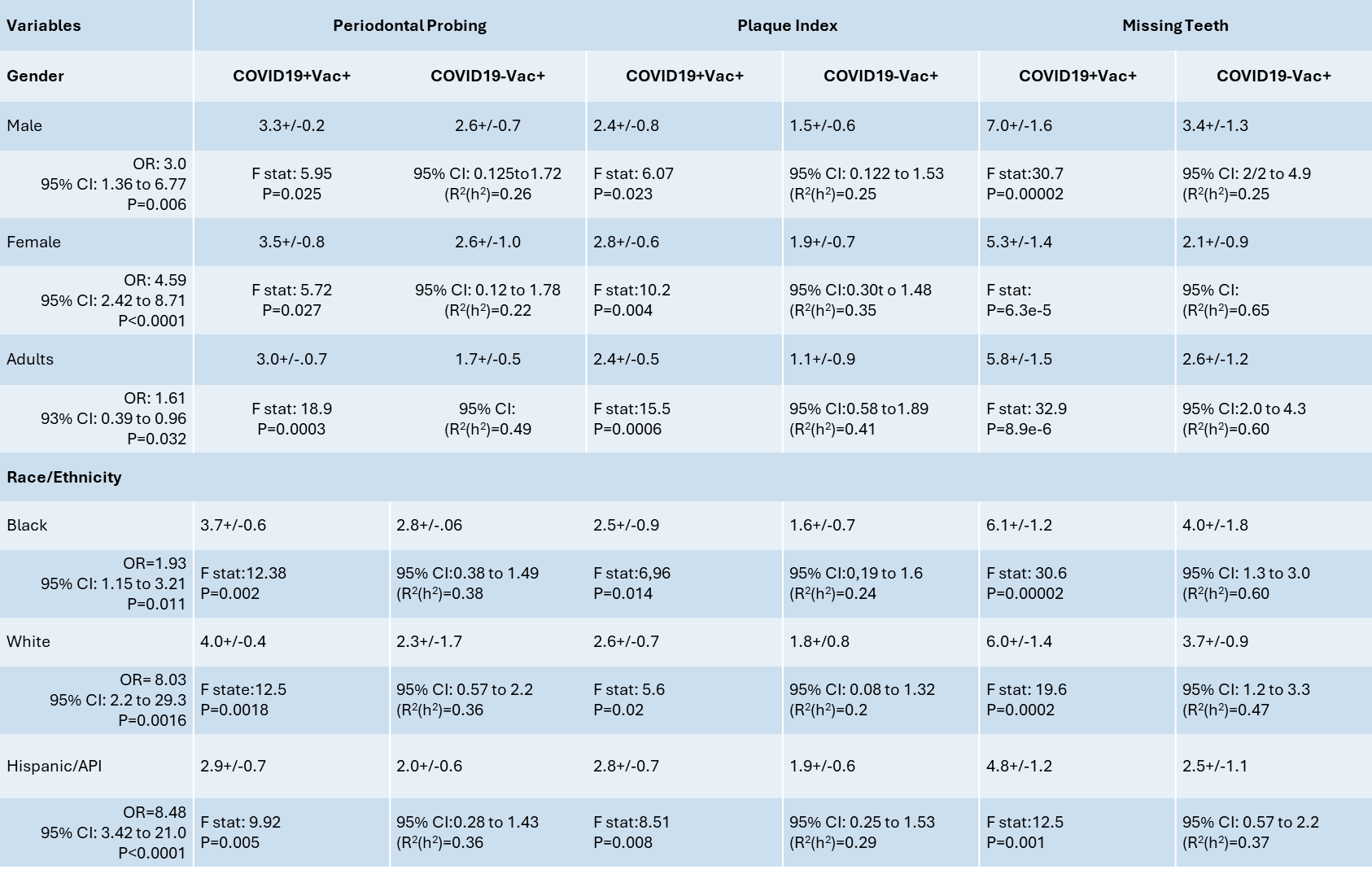

**Supplementary Table 5.** Oral Health Characteristics Related to COVID-19 Vaccination Status and History (December 2021 – March 2024; N=151) based on Gender and Race/Ethnicity Presented as Odds Ratios (OR) with 95% Confidence Intervals (CI).
